# ThX – A next-generation probe for the early detection of amyloid aggregates

**DOI:** 10.1101/713594

**Authors:** Lisa-Maria Needham, Judith Weber, Juan A. Varela, James W. B. Fyfe, Dung T. Do, Catherine K Xu, Benjamin Keenlyside, Rachel Cliffe, David Klenerman, Christopher M. Dobson, Christopher A. Hunter, Sarah E. Bohndiek, Thomas N. Snaddon, Steven F. Lee

**Author notes:** denotes joint corresponding authors. denotes equal contributions (Lisa-Maria Needham, Judith Weber).

## Abstract

Neurodegenerative diseases such as Alzheimer’s and Parkinson’s are associated with protein misfolding and aggregation. Recent studies suggest that the small, rare and heterogeneous oligomeric species, formed early on in the aggregation process, may be a source of cytotoxicity. Thioflavin T (ThT) is currently the gold-standard fluorescent probe for the study of amyloid proteins and aggregation processes. However, the poor photophysical and binding properties of ThT impairs the study of oligomers. To overcome this challenge, we have designed Thioflavin X, (ThX), a next-generation fluorescent probe which displays superior properties; including a 5-fold increase in brightness and 7-fold increase in binding affinity to amyloidogenic proteins. As an extrinsic dye, this can be used to study unique structural amyloid features both in bulk and on a single-aggregate level. Furthermore, ThX can be used as a super-resolution imaging probe in single-molecule localisation microscopy. Finally, we demonstrate that ThX can be used to detect a distinct oligomeric species, not observed via traditional ThT imaging.

## Introduction

The study of protein misfolding and aggregation is crucial for understanding the molecular mechanisms that underpin neurodegenerative diseases, such as Parkinson’s and Alzheimer’s. These neurological disorders share the hallmark of converting soluble proteins into amyloid deposits, which are characterized by a common cross-β core structural motif (Fig. 1A)^1^. Several recent studies have suggested that the small soluble oligomeric intermediates, which arise during the fibril formation process, are strongly implicated in cytotoxicity and ultimately neuronal cell death^2^. These potentially toxic oligomers are highly heterogeneous in their size and structure, and are rare in abundance (<1%) relative to the monomeric protein^3^. This renders them challenging to identify and characterize with ensemble biophysical techniques, thus there is currently a lack of suitable tools and methods to study these potentially pathogenic oligomeric species and their formation processes.

**Fig. 1.**
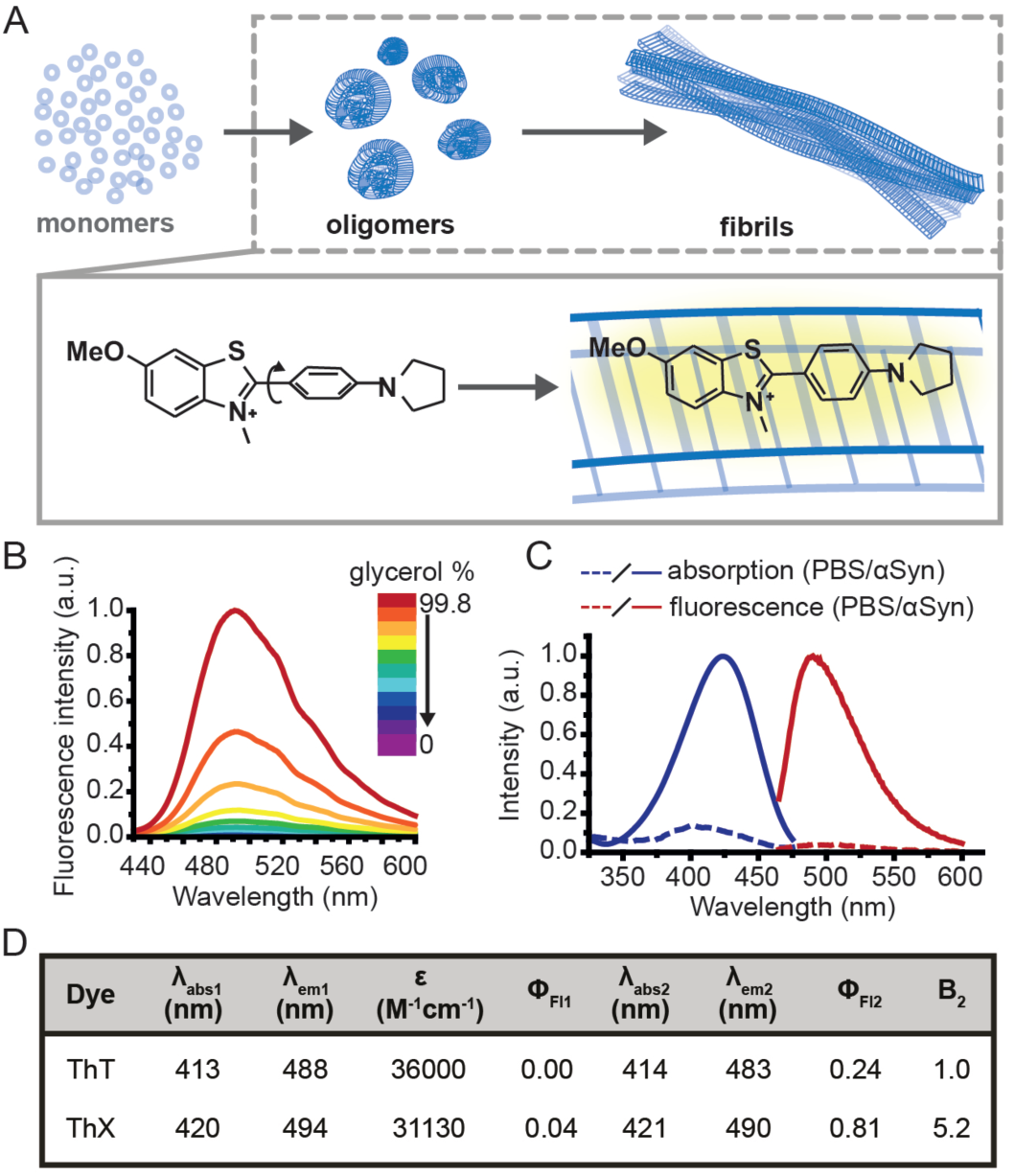
**A)** Schematic illustration of the nucleated pathway of amyloid fibril formation starting from monomeric proteins and the common cross-β sheet motif of amyloid fibrils to which benzothiazole salts bind transiently along the long fibril axis. The binding of ThX to the fibrils restricts the rotation around the carbon-carbon bond resulting in a fluorescence turn-on response.**B)** Fluorescence spectra of ThX in increasing glycerol concentration, showing the viscosity dependence of emission properties of the molecular rotor dye. **C)** Absorption and fluorescence spectra of ThX free in PBS and bound to αSyn aggregates. **D)** Table of bulk photophysical properties of ThX and ThT including; maximum absorption wavelength in PBS (λ_abs1_) and with αSyn aggregates (λ_abs2_), maximum emission wavelength in PBS (λ_em1_) and with αSyn aggregates (λ_em2_), molar extinction coefficient (ε), fluorescence quantum yield in PBS (Φ_Fl1_) and with αSyn aggregates (Φ_Fl2_) and the relative brightness (integrated fluorescence intensity) with αSyn aggregates normalized to ThT (B_2_).

The availability of improved optical probes that allow single-aggregate visualization may provide a potential solution to this problem. One of the main tools for studying amyloid fibril formation is Thioflavin T (ThT), a benzothiazole salt and molecular-rotor dye^4^, which has been used extensively as the ‘gold-standard’ in the aggregation field due to its amyloid-binding properties^5–14^. Upon binding to β-sheet-rich structures, the fluorescence intensity of ThT increases by several orders of magnitude, making it a sensitive and efficient reporter of amyloid^15^. Additionally, new microscopy techniques couple the photophysical properties of ThT with single-molecule instrumentation enabling direct observation of aggregates at diffraction-limited resolution both *in vitro*^16,17^ and in human biofluids^18,19^. Nevertheless, the low molar extinction coefficient, average quantum yield and low binding affinity of ThT mean it is poorly suited to detection of smaller oligomeric species, particularly in biological samples. Therefore, there remains an unmet need for new fluorescent probes with improved photophysical and binding properties to detect and characterize the formation of these early oligomeric species and their role in the pathogenesis of neurodegenerative disorders.

To address this, we have designed a novel ThT derivative, ThX, that outperforms ThT in its binding and optical characteristics but still retains the fluorescence enhancement attained upon binding to β-sheet containing species. To achieve this, we increased electron density on the benzothiazole core ring by exchanging the appended methyl group for the corresponding methoxy moiety, and we embedded the dimethyl amino moiety within a pyrrolidine in order to restrict rotation around the C(sp^3^)–N σ-bonds (Fig. 1A). This resulted in a more electron-rich, more conformationally restricted benzothiazole salt with increased lipophilicity. These properties improved the brightness (ε × Φ_Fl_ or integrated fluorescence intensity) upon binding to recombinant, wild-type α-Synuclein (αSyn) aggregates by 5-fold and increased the binding affinity by 7-fold. This resulted in enhanced detection of early αSyn oligomeric species with higher sensitivity both in bulk as well as at the single-aggregate level. In addition, we were able to exploit the transient nature of ThX binding to αSyn aggregates and demonstrate its excellent super-resolution imaging capabilities.

## Results and Discussion

Initially, to characterize ThX in bulk, the emission properties of the dye were assessed in aqueous solution in the absence of αSyn aggregates. ThT behaves as a molecular rotor in low viscosity environments, undergoing rapid rotation around the carbon–carbon σ-bond between the dimethylaniline and benzothiazole rings^20^. Therefore, photoexcitation induces the formation of a twisted intramolecular charge transfer (TICT) state and a low observed fluorescence quantum yield (Φ_Fl_). In high viscosity environments this rotation is more restricted resulting in an increase in Φ_Fl_. ThX exhibited viscosity dependent fluorescence intensity characteristic of ThT, which suggests that ThX also behaves as a molecular rotor (Fig. 1B). ThT is believed to bind transiently along the side chain channels that make up the β-sheet architecture of amyloids^21^ during which it is conformationally restricted (Fig. 1A). Consequently, binding induces a large increase in Φ_Fl_.

ThX also demonstrated a characteristic increase in Φ_Fl_ upon binding to αSyn aggregates in a similar manner to ThT, as a result of reduced C-C bond torsion (Fig. 1C, D, Fig. S1) However, the Φ_Fl_ of ThX in the presence of αSyn (Φ_Fl2_ = 0.81) was ∼3.4 times larger than ThT (Φ_Fl2_ = 0.24) and its brightness (defined as the total integrated emission intensity) outperformed ThT by >500%. The emission properties of ThX are a convolution of both the intrinsic photophysics and the affinity with which it binds to amyloid aggregates. Therefore, the capabilities of ThX and ThT to bind αSyn aggregates were compared. The binding affinities were obtained with direct fluorescence titrations yielding K_d_ values of 4.8 ± 1.3 μM for ThT and 0.68 ± 0.1 μM for ThX, a >700% improvement in αSyn affinity (Fig. 2Ai). This significantly increased binding affinity was mirrored at the single aggregate level and allowed ThX SAVE imaging at 3 orders of magnitude lower concentration compared to ThT, with ThX clearly obtaining high quality images of αSyn aggregates at concentrations as low as 500 pM (Fig. 2Aii, Fig. S2).

**Fig. 2.**
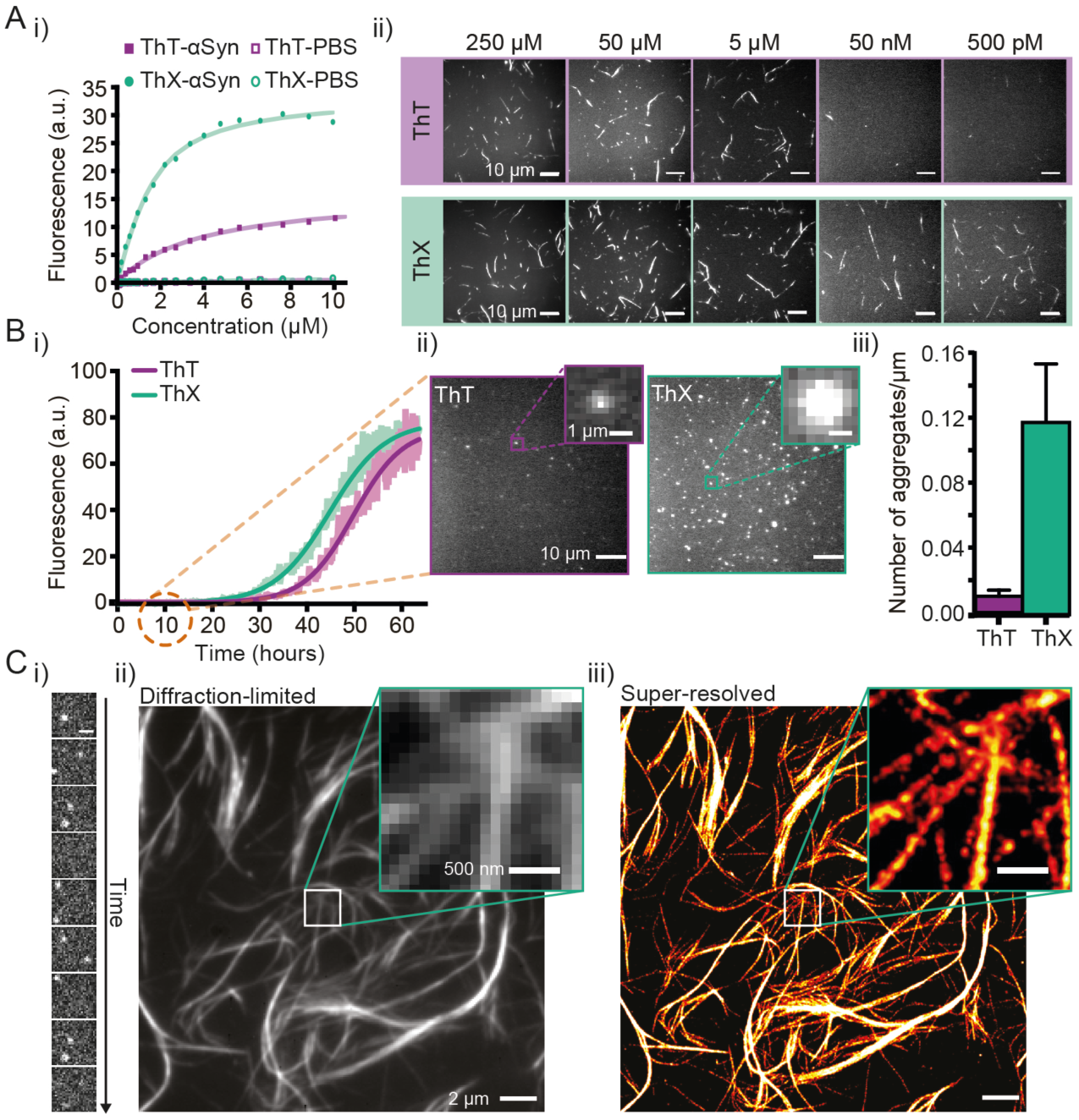
**A)** i) Representative binding curves of ThX and ThT binding to αSyn aggregates obtained by direct fluorescence titrations in the presence of 2 μM αSyn. ii) Single-aggregate fluorescence images of αSyn fibrils with ThT (top) or ThX (bottom) at concentrations between 250 μM-500 pM. **B)** i) Kinetics of αSyn amyloid fibril formation monitored by ThX and ThT. The solid lines indicate the fitted sigmoidal growth curve, the brighter shadows the averaged values obtained from a minimum N=7 independent αSyn aggregations. ii) Fluorescence images taken with 5 μM ThT (left) and ThX (right) of single αSyn aggregates formed 10 hours into the aggregation. The insets show a representative single aggregate. iii) A bar graph showing the density of single 10-hour αSyn aggregates detected with ThT and ThX. Error bars represent standard deviations from 27 fields of view. **C)** i) Time montage showing single ThX molecules transiently binding to a region of an αSyn fibril (scale bar = 500 nm.) ii) Diffraction limited and ii) super-resolved image of αSyn fibrils. Detailed structural features are obscured in the diffraction limited image which can only be seen once the image has been super-resolved (insets).

Next, we explored whether ThX can be used to monitor the process of amyloid formation through protein aggregation. First, using ThT and ThX, bulk kinetic measurements of recombinant αSyn protein aggregations were performed in parallel. The formation of αSyn fibrils followed the expected sigmoidal growth curves, consistent with multiple previous studies^12,14,22^(Fig. 2Bi). A clear signal with an intensity of 5% above the baseline signal, was reached at 26.3 ± 1.4 hours for ThX and 34.1 ± 0.8 hours for ThT (Fig. S3). These data indicated that ThX was able to detect aggregate formation at significantly earlier times than ThT in the bulk. This was further confirmed at the single aggregate level with ThX detecting 7.5-fold more species than ThT 10 hours into the aggregation (Fig. 2Bii). The differences were not caused by different abilities of ThX and ThT to inhibit the aggregation process (Fig. S4). In addition, ThX a generic probe and also outperforms ThT in its detection capabilities with other amyloid proteins such as amyloid-β peptide (Aβ_1-42_) and P301S tau at both the bulk (Fig. S5) and single aggregate level (Fig. S6).

Although ThX demonstrated superior photophysical and binding properties in the bulk and single-aggregate detection regimes, smaller spatial information, (i.e. oligomeric species) are still obscured by the diffraction limit (∼250 nm). Spehar and Lew *et al.* previously demonstrated the use of ThT as a super-resolution probe by exploiting the transient nature of its binding to amyloid proteins and its low solution Φ_Fl_ in a method termed transient amyloid binding (TAB)^23^ Interestingly, ThX obeys similar principles; by excitation with 488 nm light and collection at ∼587 nm, one can generate isolated fluorescent puncta of single ThX molecules (Fig 2Ci, Supplementary movie 1). These localisations can be summed to generate diffraction-limited images of αSyn fibrils (Fig. 2Cii) as well as fit and reconstructed to produce super-resolved images (Fig. 2Ciii) of αSyn fibrils, achieving a mean localisation precision of 21.0 nm (Fig. S7A) and an image resolution of 18.3 nm (Fig. S7B), determined by Fourier ring correlation analysis^24^.

Amyloid aggregates are diverse in size, structure and formation mechanism. Previous work has provided evidence for a structural rearrangement from disordered aggregates to ordered fibrils during the aggregation process^19,25,26^. We have previously developed an all-optical method to characterize the structural order of individual protein aggregates by measuring the fluorescence anisotropy of bound ThT^19^. We employed this technique to elucidate whether ThX was sensitive to alternative binding modes in both αSyn fibrils and oligomers. For both ThX and ThT the fluorescence intensity modulated sinusoidally in the same phase when bound to ordered structures (Fig. S8), suggesting that the two dyes are oriented the same way in this circumstance. However, when applied to early αSyn aggregates (6-hour time point, Fig. 3A) ThX could observe large numbers of both modulating (>0.5, β-sheet ‘ordered’) and non-modulating (<0.5, β-sheet ‘disordered’) distributions (Fig. 3Bii). This disordered aggregate population was not detected when using ThT alone (Fig. 3Bi). Photon-fluence matched experiments confirmed that both species (albeit fewer in number) could be observed with ThX but not ThT (Fig. S9) and that this was not just a product of the increased affinity. Furthermore, the degree of modulation was not correlated to the fluorescence intensity (Fig. 3Ci-ii, Fig S10). This may suggest that the number of binding sites remains unchanged during the structural rearrangement of the aggregates, and that the ability of ThX to detect different species is not just a result of the increased brightness of8ThX. In combination, these data support the notion that ThX is able to detect a structurally distinct oligomeric species. We expect that the capabilities of this new probe will be of broad interest to the aggregation community.

**Fig. 3.**
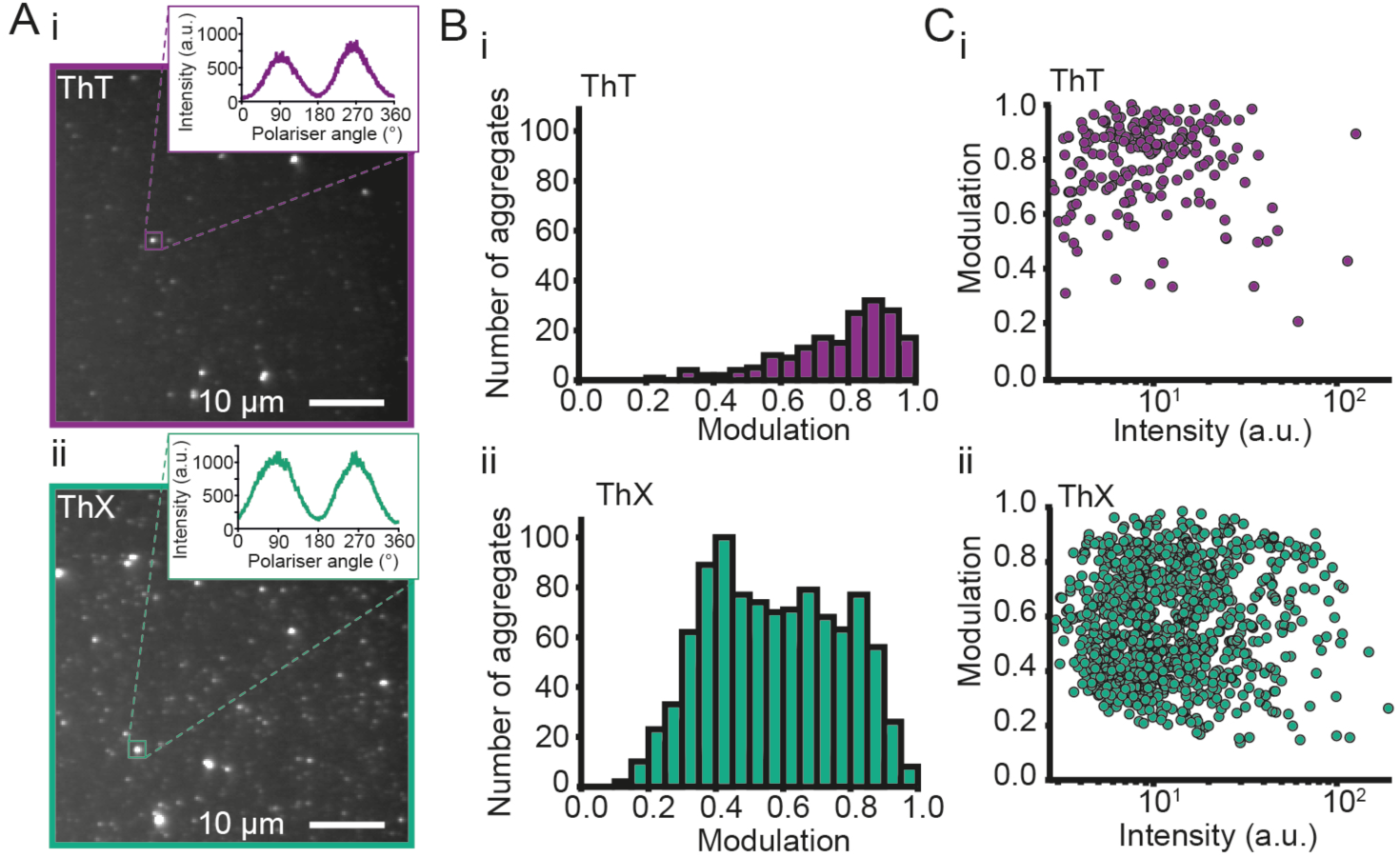
**A)** Representative fluorescence anisotropy images of αSyn aggregates 6 hours into the aggregation reaction with i) ThT and ii) ThX. **B)** Histograms of the extent of modulation of single αSyn aggregates detected with i) ThT and ii) ThX. **C)** Relationship between extent of modulation of αSyn aggregates and i) ThT or ii) ThX mean fluorescence intensity measured during the rotation of the polariser minus the background intensity. For each dye, N=3 separate aggregation reactions were tested, and slides were imaged at minimum three different fields of view.

## Conclusions

We have designed and evaluated ThX, a novel ThT derivative with improved photophysical and binding properties that may prove superior to ThT in future applications. ThX can detect β-sheet species at early time points in the aggregation of amyloid proteins at both the bulk and single-aggregate levels, as well as additional structurally distinct species that are not observed with ThT. In addition, ThX is compatible with super-resolution imaging and capable of resolving nano-scale structural features with ∼20 nm precision. Taken together, ThX may allow the study of the formation of earlier pathological amyloid species and exploration of their role in the pathogenesis of neurodegenerative disorders. Furthermore, the unique properties of ThX might enable the study of these amyloids in high background biological samples such as cerebrospinal fluid and blood plasma.

## Contributions

J.W., L.M.N., S.F.L. and T.N.S. designed the experiments, T.N.S., J.W.B.F. and D.T.D. synthesized the molecules. J.W. and L.M.N. performed bulk fluorescence characterisation measurements and analyses. J.W. prepared the αSyn aggregates, performed bulk protein binding assays and aggregation time-course experiments. J.W. and C.A.H. analysed binding data. L.M.N. performed the single-aggregate fluorescence imaging, super-resolution imaging and analyses. J.W. and J.A.V. performed fluorescence polarisation experiments. J.A.V. analysed fluorescence polarisation data. J.W., L.M.N., S.F.L. and T.N.S. wrote the manuscript.

### Acknowledgements

We thank the Royal Society for the University Research Fellowship of S.F.L. (UF120277). This work was funded in part by the Michael J Fox Foundation and The University of Indiana. We thank the EPSRC for the Doctoral Prize of L.M.N. J.W, and S.E.B. are funded by Cancer Research UK (C14303/A17197, C47594/A16267), the EPSRC-CRUK Cancer Imaging Centre in Cambridge and Manchester (C197/A16465) and the European Union’s Seventh Framework Programme (FP7-PEOPLE-2013-CIG-630729).

We thank Ewa Klimont and Swapan Preet for α-synuclein protein expression.

We thank Jane Gray and Ian Hall from the research instrument core facility of the Cancer Research UK Cambridge Institute for technical support.

