## Supplementary Information and Methods for "ThX – A next-generation probe for the early detection of amyloid aggregates"

* - denotes joint corresponding authors

$ - denotes equal contributions

### Contents

Supplementary Results .......................................................................................................1

Experimental Methods .......................................................................................................6

### Supplementary Results

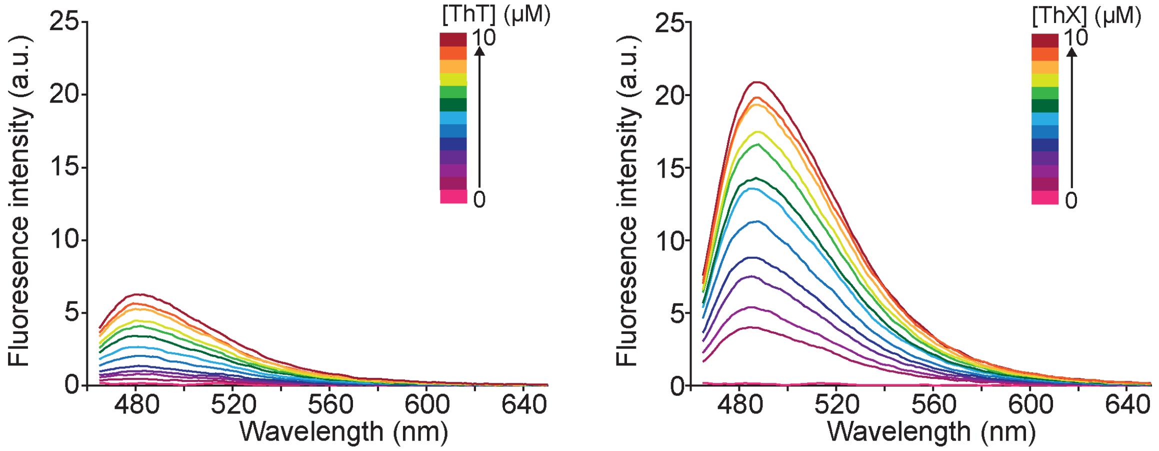


**Figure S1.** Fluorescence spectra of ThT (left) and ThX (right) at concentrations between 0 and 10 μM in the presence of 2 μM αSyn aggregates.

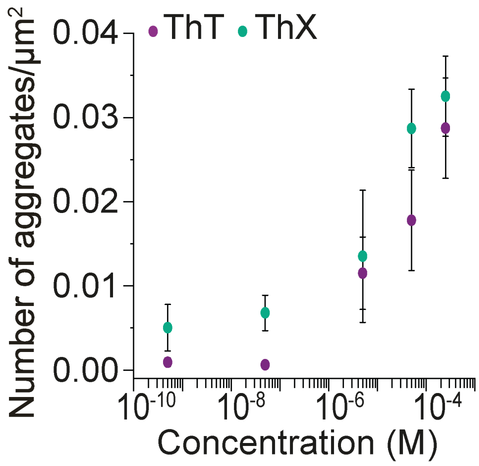


**Figure S2.** Number of single αSyn aggregates detected on a surface during a single-aggregate imaging experiment with varying ThT/ThX concentrations. Error bars represent the standard deviations of aggregates across 27 fields of view.

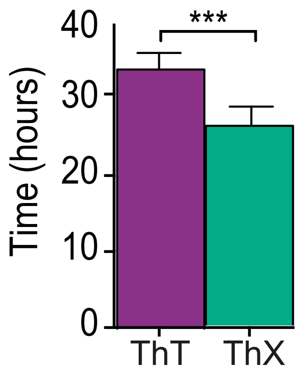


**Figure S3.** Bar graph indicating the time-points in the αSyn aggregation curve when the fluorescence signal from ThT/ThX was 5% above the baseline signal. Error bars are standard deviations from a minimum of 7 independent αSyn aggregations.

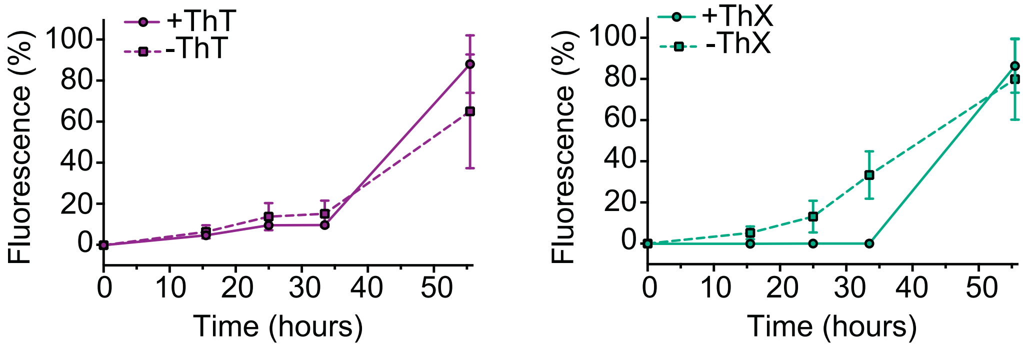


**Figure S4.** Graphs showing the ThT (left) and ThX (right) fluorescence during an αSyn aggregation time-course with ThT/ThX either present (solid line) or absent (dashed line) from the reaction.

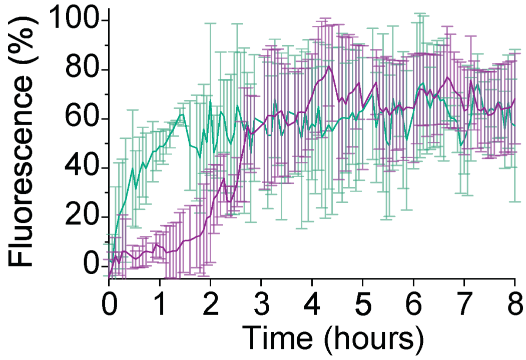


**Figure S5.** Graph showing the ThT/ThX fluorescence during an A$\beta_{1\text{-}42}$ aggregation time-course. The A$\beta_{1\text{-}42}$ protein is known to exhibit a fast-kinetic profile in comparison to αSyn, with a short lag phase between 1 - 2 hours when measured in bulk with ThT. The kinetic curve obtained with ThT was in accordance with the literature and did reveal a lag phase of ~1 hour with a plateau reached at ~6 hours. In comparison to this, nearly no lag phase was observed with ThX and the plateau phase was reached after 3 - 3.5 hours. Error bars represent the standard deviations from 3 aggregations

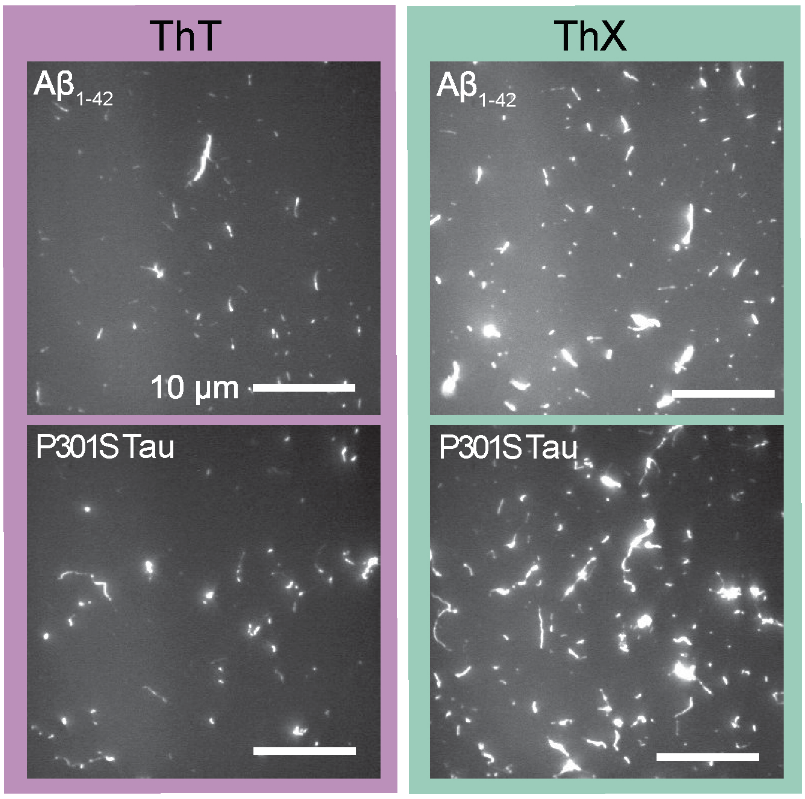


**Figure S6.** Single-aggregate fluorescence images of A$\beta_{1-42}$ peptide (top) and P301S tau (bottom) taken with ThT (left) and ThX (right). ThX was able to detect 1.7-fold and 1.8-fold more A$\beta_{1-42}$ and P301S tau aggregates respectively compared to ThT.

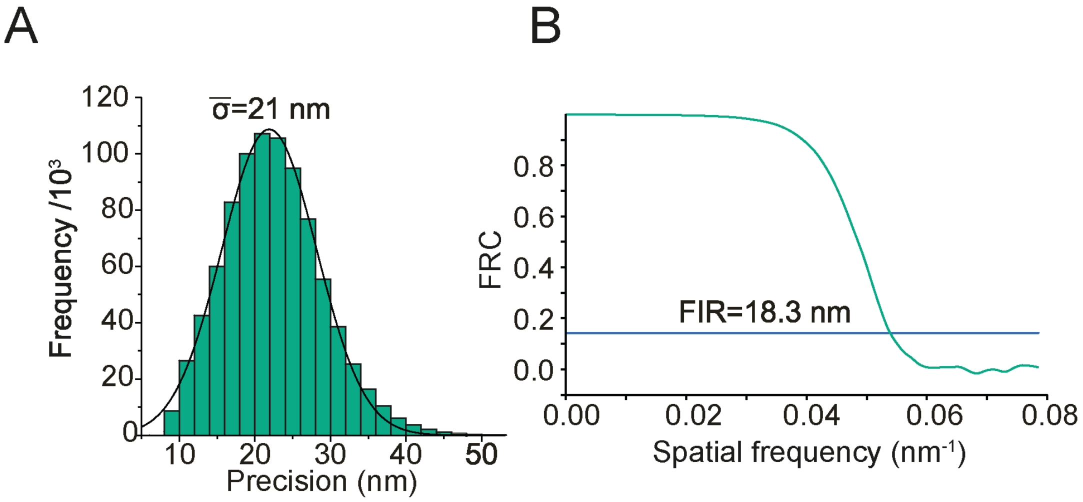


**Figure S7. A)** Histogram of localisation precision of single ThX emitters. B) Fourier ring correlation analysis showing the Fourier image resolution (FIR) of the ThX super-resolved image of αSyn fibrils.

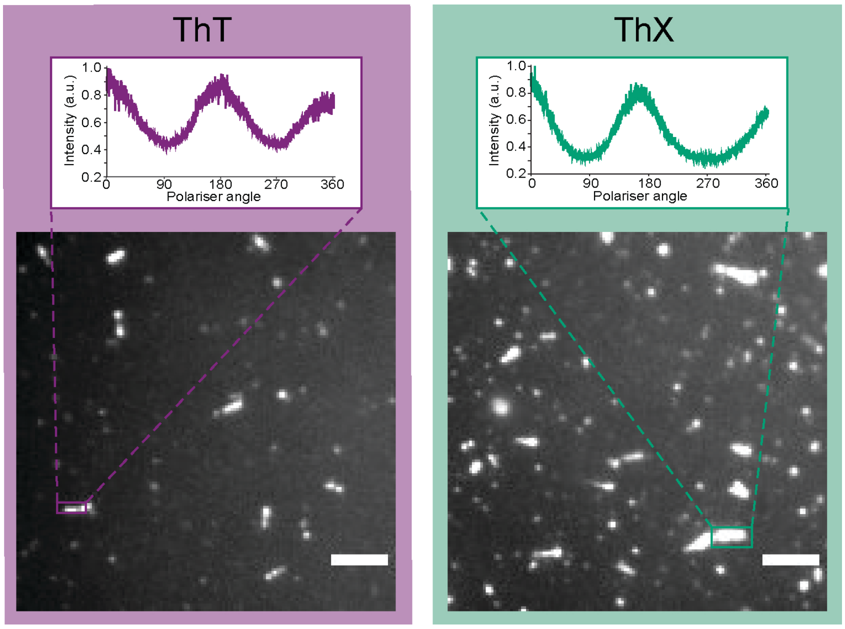


**Figure S8.** - Representative fluorescence anisotropy images of late stage (>72 hours) αSyn aggregates with ThT (left) and ThX (right). Scale bar= 5 μm.

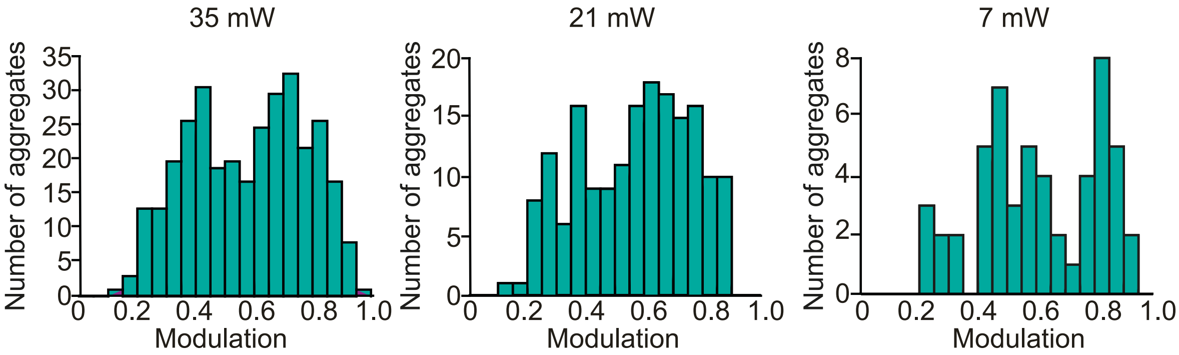


**Figure S9.** Histograms of the modulation of values of αSyn aggregates (6-hours) measured in fluorescence polarisation experiments with ThX using different laser powers; 35mW (left), 21mW (middle) and 7 mW (right).

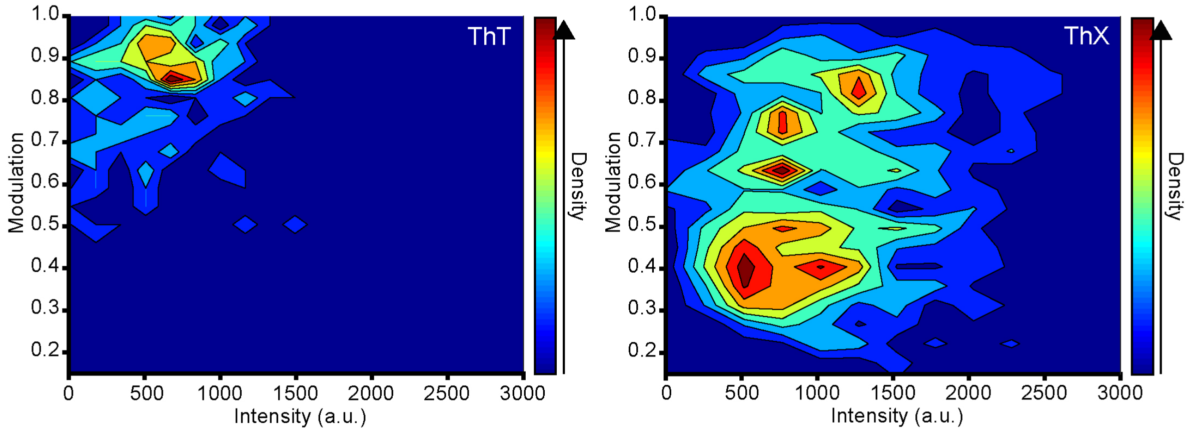


**Figure S10**. Density plots illustrating the relationship between extent of modulation of αSyn aggregates and ThT (left) or ThX (right) mean fluorescence intensity measured during the rotation of the polariser minus the background intensity. These data are presented as scatter plots in Figure 3Ci-ii.

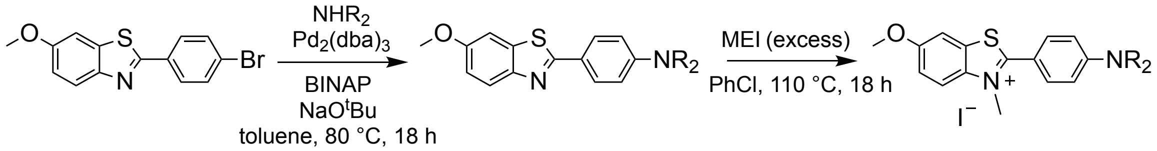


**Figure S11.** Synthetic route from a bromide precursor to ThX.

**Supplementary movie 1-** Raw movie of single-molecules of ThX (1 μM) transiently binding to αSyn fibrils on a surface. ThX was excited with 488 nm laser light in a TIRF geometry. Gold beads (100 nm) were used as fiducial markers to account for nanoscale XY drift. The scale bar is 5 μm.

### Experimental Methods

#### Synthesis of ThX

**General Considerations.** All reactions (Fig. S11) were magnetically stirred, and solvents purified under a dry argon atmosphere by passage through activated alumina and Q5 Grubbs apparatus. Workup and purification procedures were conducted in air using reagent grade solvents purchased from Sigma-Aldrich. Reactions were monitored by thin layer chromatography (TLC) using SiO2 on pre-coated glass plates, and a layer thickness of 0.25 mm. Compounds were visualized by UV light (254 and 366 nm) and a basic potassium permanganate stain (3 g KMnO4, 20g K2CO3, 2.5 mL 10% aq. NaOH, in 400 mL DI H2O). Column chromatography was performed with silica gel 60 (35-70 µm). IR spectra were recorded on an Avatar 360-FT IR E.S.P. on a NaCl salt plate. Absorptions are reported as values in cm-1 followed by the relative intensity: s = strong, m = medium, and w = weak. 1H NMR spectra were recorded at room temperature on a Varian I400, (400 MHz), I500 (500 MHz), I600 (600 MHz) or a Varian VXR400 (400 MHz) Fourier Transform spectrometer using an internal deuterium lock. The chemical shift in ppm is quoted relative to the residual solvent peak of chloroform (∂H = 7.26, ∂C = 77.05) as the internal standard, unless otherwise reported. Multiplicities are described as s = singlet, d = doublet, t = triplet, m = multiplet. Coupling constants are reported in Hz, and integration. Low mass spectrometry was carried out on an Agilent 1200 HPLC-6130 MSDAll using ESI and mass to charge ratios (m/z) are reported as values in atomic mass units.

6-methoxy-2-(4-(pyrrolidin-1-yl)phenyl)benzo[d]thiazole[1] (S1):

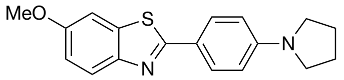


An oven-dried microwave vial was charged with 2-(4-bromophenyl)-6-methoxybenzo[d]thiazole2 (480 mg, 1.5 mmol, 1 equiv), Pd2(dba)3 (1 mol%), ( ±)-BINAP (3 mol%) and NaOtBu (1.4 equiv) in nitrogen-filled glovebox. Pyrrolidine (1.2 equiv.), was then added and the vial was capped with microwave vial cap. PhCH3 (3 mL) was added and the resulting solution heated to 80 °C for 18 hours. The reaction cooled to room temperature, diluted with EtOAc (5 mL) and filtered through Celite. The filtrate was concentrated, and the residue was triturated with Et2O (20 mL). The solid was filtered and rinsed with cold $Et_2O$ to give *6-methoxy-2-(4-(pyrrolidin-1-yl)phenyl)benzo[d]thiazole* (65 mg, 0.9 mmol, 60%) as a colorless solid.

Melting point: 215–216 °C

IR (Film): 2967.53, 2845.34, 1605.96, 1552.67, 1462.86, 1427.33, 1318.03, 1263.69, 1225.98, 1183.24, 1028.35, 1005.23, 969.51, 860.75 $cm^{-1}$.

$^{1}H$ NMR (400 MHz, $CDCl_3$) δ = 7.89 (d, *J* = 8.8 Hz, 2H), 7.85 (d, *J* = 8.9 Hz, 1H), 7.31 (d, *J* = 2.6 Hz, 1H), 7.03 (dd, *J* = 8.9, 2.6 Hz, 1H), 6.59 (d, *J* = 8.8 Hz, 2H), 3.88 (s, 3H), 3.37 (m, 4H), 2.05 (m, 4H).

$^{13}C$ NMR (125 MHz, CDCl3): δ = 160.2, 150.5, 142.9, 142.4, 129.2, 122.1, 116.1, 114.4, 108.2, 49.3, 41.0, 18.9.

HRMS (ESI): m/z calculated for [M+H] $C_{18}H_{19}N_2OS$: 311.1128. Found: 311.1217

6-methoxy-3-methyl-2-(4-(pyrrolidin-1-yl)phenyl)benzo[d]thiazol-3-ium iodide:

Bromide S1 (62 mg, 0.2 mmol, 1.0 equiv) and methyl iodide (1mL, excess) were added to an oven-dried 4 mL vial containing a magnetic stir bar equipped with a Teflon septa insert screw cap. Chlorobenzene (1.5 mL) was added via syringe and the reaction heated to 110 ºC as stirred for 18 h. The reaction was then cooled to room temperature and directly triturated with $Et_2O$. The colorless solid was collected by filtration to give the *6-methoxy-3-methyl-2-(4-(pyrrolidin-1-yl)phenyl)benzo[d]thiazol-3-ium iodide* (81 mg, 0.18 mmol, 90%).

Melting point: 208–210 °C

IR (film): 2972.42, 2864.46, 1601.64, 1536.63, 1503.98, 1481.67, 1441.1, 1298.23, 1272.35, 1064.88, 1033.87, 1015.83, 954.81, 859.25 $cm^{-1}$.

$^{1}H$ NMR (500 MHz, $CDCl_3$): δ = 7.94 (dd, *J* = 9.3, 1.8 Hz, 1H), 7.89 (d, *J* = 2.2 Hz, 1H), 7.35 – 7.28 (m, 1H), 6.71 – 6.64 (m, 2H), 4.43 (d, *J* = 1.8 Hz, 3H), 3.90 (d, *J* = 1.8 Hz, 3H), 3.4 (m, 4H), 2.08 (m, 4H),

$^{13}C$ NMR (126 MHz, $CDCl_3$) δ 170.9, 159.4, 151.5, 136.7, 132.6, 129.7, 129.7, 128.6, 126.4, 118.9, 117.4, 112.6, 110.4, 106.7, 56.7, 47.9, 39.9, 25.4.HRMS (CI): m/z calculated for [M–I] $C_{19}H_{21}N_2OS$: 325.1375. Found 325.1382.

**Preparation of αSyn aggregates.** The purification of monomeric wild-type αSyn from Escherichia coli was performed following a published protocol[3]. αSyn aggregates used for bulk photophysical characterization and binding affinity measurements were generated as previously described[4]. Briefly, monomeric αSyn was diluted into 50 mM sodium phosphate (pH 7.4) to reach a final concentration between 150 and 300 μM. Samples were incubated at 45°C for in total 72 hours whilst stirring at maximum speed with a Teflon flea. The aggregation reaction was sonicated at 24-hour intervals (3 cycles of 10 seconds at 30% power (Branson digital Sonifer, model 250, Branson, St Louis MO, USA). Finally, the aggregation mixture was aliquoted, flash frozen in liquid $N_2$ and stored at -80 °C. The αSyn concentration stated in the different experiments is reported as the monomer equivalent concentration.

For all single-aggregate and super-resolution imaging experiments, αSyn aggregates were generated by diluting monomeric αSyn into filtered (0.2 μm syringe filter, Whatman, 6780-1302) PBS buffer containing 0.01% $NaN_3$ to a final concentration of 70 μM. The solution was then incubated in the dark at 37 °C with constant shaking (200 rpm) for the time stated in the experiment.

**Preparation of ThX and ThT stock solutions.** Stock solutions of ThX and ThT (Sigma-Aldrich, T3516) were prepared by dissolving the solid into dimethyl sulfoxide (DMSO, Sigma Aldrich, 276855) to a concentration of 10 mM, these were stored in the dark at -80 °C. The stock solutions were diluted into filtered (0.02 μm syringe filter, Whatman, 6809-1102) PBS (pH 7.4) to a concentration of 500 μM. The diluted dye solutions were then filtered (0.02 μm syringe filter, Whatman, 6809-1102). The diluted solutions were stored in the dark at 4 °C for a maximum of a week after preparation.

**Photophysical characterization.** Samples were placed into a quartz fluorescence cuvette (Hellma Analytics, 3 × 3 mm) and bulk fluorescence spectra were recorded using a fluorescence spectrophotometer (Cary Eclipse, Varian) in which emission was collected perpendicular to direction of propagating excitation. Initial fluorescence spectra of ThX at multiple concentrations (0.25-25 μM) were recorded free in PBS (pH 7.4) and in the presence of late stage αSyn aggregates (~2 μM). Replicates were made with n=3 separately prepared dye samples. UV-Vis absorption spectra were collected by placing samples in a quartz transmission cuvette (Hellma Analytics, 1 cm path length) and measured with an absorption spectrometer (Cary 400, Varian). Buffer blanks were collected for both fluorescence and absorption spectra and used for background correction. Viscosity dependent fluorescence measurements of ThX (10 μM) were recorded in PBS containing glycerol (0-99.8%, Sigma-

Aldrich, G9012). Several photophysical quantities of ThX and ThT were determined from ensemble absorption and emission data using a custom Matlab script. These included absorption maxima ($\lambda_{abs}$), emission maxima ($\lambda_{em}$), molar extinction coefficient ($\varepsilon$), fluorescence quantum yield ($\Phi_{Fl}$) and brightness (B). $\varepsilon$ was determined from a calibration curve of ThX/ThT of the UV-vis absorbance and sample path length using the Beer–Lambert law with solutions of known concentrations. $\Phi_{Fl}$ of the dyes were determined by referencing against rhodamine 101 (Sigma-Aldrich, 83694) in ethanol ($\Phi_{Fl}$= 1.0) and discrepancies in absorbance and solvent refractive index were corrected for[5]. Fluorescence emission spectra of rhodamine 101 were collected at several concentrations (100 pM- 100 μM).

**Fluorescence-based binding assay.** All samples were prepared in low-binding Eppendorf tubes (Eppendorf AG, Hamburg, Germany). Fluorescence and absorption titrations were recorded using a fluorescence spectrophotometer (Cary Eclipse, Varian) and a UV-VIS spectrophotometer (Cary 300 Bio, Varian). αSyn aggregates (4 μM) were prepared in filtered (0.02 μm syringe filter, Whatman, 6809-1102) PBS. Dye stock solutions were diluted into filtered PBS to prepare seven solutions with concentrations ranging between 19.5 μM and 1 mM, which were then sonicated carefully. The 4 μM αSyn dilution was diluted to 2 μM, transferred into a quartz cuvette (Hellma Analytics, 3 × 3 mm) and the fluorescence and absorption spectra were recorded. Defined volumes of dye dilutions and the same volume of 4 μM αSyn dilutions - to avoid dilution of αSyn during the titration - were successively added to the cuvette. The solution was thoroughly mixed, and the spectra were recorded after each addition. The same protocol was carried out without the presence of αSyn. Fluorescence spectra were acquired with an excitation wavelength of 440 nm and emission was collected between 465 nm and 600 nm with an emission slit of 5 nm. Fluorescence spectra were analyzed using a Microsoft Excel spreadsheet to fit the changes in fluorescence intensity at fixed wavelengths to a 1:1 binding isotherm by optimizing the association constant and fluorescence intensity of the free and bound dye using bespoke VBA macros (equation).

**Measuring aggregation kinetics of wild-type αSyn.** To study the aggregation kinetics of αSyn, wild-type αSyn monomers were diluted into filtered (0.2 μm syringe filter, Whatman, 6780-1302) PBS (pH 7.4) containing $NaN_3$ (0.01%) and ThT or ThX (18 μM) to reach a final concentration of 70 μM. Each sample was then pipetted into multiple wells of a 96-well, low-binding polyethylene glycol coating plate (Corning 3881, Kennebuck ME, USA) with a clear bottom, at 100 μL per well. The 96-well plate was sealed with a gas permeable plate seal, placed in a plate reader (CLARIOstar; BMG Labtech, Ortenberg, Germany) and incubated at 37°C. Fluorescence was measured using excitation and emission filters of 440-10 nm and

490-15 nm. Measurements were performed every 10 min for a period of up to 63.5 hours. Plates were subjected double orbital shaking at 200 rpm for 540 seconds after each reading.

**Slide preparation for fluorescence imaging.** Glass cover-slides (VWR, 631-1570) for single-aggregate fluorescence imaging were cleaned for 1 hour using an argon plasma (PDC-002, Harrick Plasma). Frame-Seal slide chambers (9 x 9 mm, Bio-Rad, Hercules,CA, product number SLF-0201) were attached to the cover-slide and the surface incubated with poly-l-lysine (50 µL, PLK, Sigma-Aldrich, 0.1% w/v, P820) for 30 minutes. The slide was then washed three times with filtered (0.2 µm syringe filter, Whatman, 6780-1302) PBS prior to imaging.

**Microscope set-up for single-aggregate imaging and fluorescence polarisation experiments.** Imaging was performed on a bespoke total internal reflection fluorescence microscope using a 405 nm excitation laser (LBX-405-50-CIR-PP, Oxxius), which was passed through a quarter wave plate (WPQ05M-405, Thorlabs) aligned parallel to the optical axis at the edge of an objective lens (60x Plan Apo TIRF, NA 1.45 oil-immersion, Nikon Corporation), mounted on an inverted optical microscope (Eclipse TE2000-U, Nikon Corporation). Fluorescence emission was also collected by the same objective and selected by the presence of a dichroic (ZT 405/532 rpc) and subsequently passed through an emission filter (FF01-480/40/25) and focused by a tube lens onto an EMCCD camera (Evolve 512, Photometrics). Micromanager software was used to control the microscope[6].

For fluorescence polarisation experiments, a polariser (10LP-VIS-B, Newport) mounted on a stepper motor rotation mount (K10CR1/M, Thorlabs) was placed between tube lens and the microscope body (described above). During image acquisition, the polariser was rotated 360 degrees at 2 degrees per second.

**Microscope set-up for super-resolution imaging.** Imaging was performed on a bespoke total internal reflection fluorescence microscope using a 488 nm excitation laser (iBeam-SMART, Toptica). The beam was circularly polarized using quarter-wave plates specific to each wavelength and then expanded, collimated and aligned parallel to the optical axis at the edge of an objective lens (100x Plan Apo TIRF, NA 1.49 oil-immersion, Nikon Corporation) mounted on an inverted optical microscope (Ti-E, Eclipse, Nikon Corporation). Fluorescence emission was collected by the same objective lens and separated from excitation light using a dichroic mirror (Di01-R405/488/561/635, Semrock) and passed through appropriate emission filters (LP02-568RS-25, FF01-587/35-25, Semrock). The fluorescence was then expanded and focused on an electron-multiplying charge-coupled device (Evolve 512, Photometrics) for imaging. Each pixel on the final image was equal to 98.6 nm.

**Single-aggregate fluorescence imaging.** αSyn fibrils (72-hour, 70 µM) were diluted into filtered (0.2 µm syringe filter, Whatman, 6780-1302) PBS in low-binding tubes (Eppendorf AG, Hamburg, Germany) to a final concentration of 1 µM prior to imaging. Pre-diluted solutions of ThT/ThX (500 µM) were further diluted to the desired imaging concentration (500 pM-250 µM for concentration dependence, 5 µM for all other imaging experiments). The diluted fibrils (50 µL) were incubated on clean, PLK coated glass cover-slides for 10 minutes. The solution was then pipetted up and down vigorously to maximise adherence of the protein to the surface following which the slide was washed three times with filtered PBS. A solution of ThT/ThX (50 µL) was then added and the slide imaged immediately. 200-frame image stacks were recorded on the microscope set-up described above in frame-transfer mode at 20 frames per second with an exposure time of 50 ms. Each pixel was equal to 241 nm. Three cover-slides and 27 separate fields of view per slide were collected for each condition. To correct for any fluorescent artefacts caused by the cover-slides or any of the solutions being used, dye solutions (5 µM, 50 µL) in PBS without αSyn were imaged for each experiment and used as blanks.

**Fluorescence polarisation measurements.** Polarisation with αSyn fibrils: αSyn aggregates (48 hours 70 µM [monomer concentration]) were diluted into filtered (0.2 µm syringe filter, Whatman, 6780-1302) PBS to a final concentration of 7 µM. Pre-diluted solutions of ThT/ThX (500 µM) were further diluted to the desired imaging concentration (5 µM). The diluted fibrils (50 µL) were incubated on clean, PLK coated glass cover-slides for 10 minutes. The solution was then pipetted up and down vigorously to maximise adherence of the protein to the surface following which the slide was washed three times with filtered PBS. A solution of ThT/ThX (50 µL) was then added and the slide imaged immediately. For each condition a minimum of 2 different cover-slides were imaged at 3-5 different fields of view per cover-slide.

Polarisation with early-stage αSyn aggregates: αSyn aggregates (6 hours, 70 µM) were diluted into filtered (0.2 µm syringe filter, Whatman, 6780-1302) PBS to a final concentration of 7 µM. Pre-diluted solutions of ThT/ThX (500 µM) were further diluted to the desired imaging concentration (5 µM). The diluted fibrils (50 µL) were incubated on clean, PLK coated glass cover-slides for 10 minutes. The solution was then pipetted up and down vigorously to maximise adherence of the protein to the surface following which the slide was washed three times with filtered PBS. A solution of ThT/ThX (50 µL) was then added and the slide imaged immediately. For each dye, three separate aggregation reactions were tested, and slides were imaged at minimum three different fields of view.

3000-frame image stacks were recorded on the microscope set-up described above in frame-transfer mode at 20 frames per second with an exposure time of 50 ms. Each pixel was equal to 241 nm. The image stacks corresponding to the full spin of the polariser were denoised in ImageJ (NIH, Bethesda) by sub-averaging 30 frames. To correct for the small circular drift caused by the rotation of the polariser, the images were aligned using “Align Images” plug-in available in GDSC utility plug-ins from the University of Sussex (http://www.sussex.ac.uk/gdsc/intranet/ microscopy/imagej/utility). After alignment, the entire stack was averaged and by using the Find Maxima function in ImageJ each protein aggregate was detected. The threshold was set to 50. To obtain the modulation signal for each species detected, analysis was performed as described in the literature[7]. Shortly, the signal intensity of a 3×3 pixels square in each detected species was calculates and the local background was subtracted. The resulting signal was then plotted and fitted with a sinusoidal curve of the form $y = a\cos(bx+c)+d$. In case of the phase c of the aggregate being similar to the phase c of the background, the aggregate was not considered in the analysis. If the quality of the sinusoidal fit was appropriate, modulation was calculated by modulation = 2a/(a+d).

**Super-resolution imaging.** αSyn fibrils (72-hour, 70 μM) were diluted into filtered (0.2 μm syringe filter, Whatman, 6780-1302) PBS in low-binding tubes (Eppendorf AG, Hamburg, Germany) to a final concentration of 2 μM prior to imaging. Pre-diluted solutions of ThX (500 μM) were further diluted to the desired imaging concentration (1 μM). Gold nanoparticles in PBS (100 nm, 753688, Sigma-Aldrich) were used as fiducial markers and were diluted to 2 μM and incubated on clean, PLK coated glass cover-slides for 5 minutes. The slide was then washed three times with filtered PBS. The diluted fibrils (50 μL) were then incubated on the cover-slide for 10 minutes. Following this, the solution was then pipetted up and down vigorously to maximise adherence of the protein to the surface following which the slide was washed three times with filtered PBS. A solution of ThX (50 μL) in PBS was then added and the slide imaged immediately. 20,000-frame image stacks were recorded on the microscope set-up described above in frame-transfer mode at 50 frames per second with an exposure time of 20 ms. To correct for any fluorescent artefacts caused by the cover-slides or any of the solutions being used, dye solutions (1 μM, 50 μL) in PBS without αSyn were imaged for each experiment and used as blanks. Super-resolution images were reconstructed using the Drift Calculator and Peak Fit package (GDSC SMLM, University of Sussex) in ImageJ.

3 W. Hoyer, T. Antony, D. Cherny, G. Heim, T. M. Jovin and V. Subramaniam, *J. Mol.*

*Biol.*, 2002, **322**, 383–393.

4 A. K. Buell, C. Galvagnion, R. Gaspar, E. Sparr, M. Vendruscolo, T. P. J. Knowles, S. Linse and C. M. Dobson, *Proc. Natl. Acad. Sci.*, 2014, **111**, 7671–7676.

5 J. R. Lakowicz, *Principles of Fluorescence Spectroscopy Principles of Fluorescence Spectroscopy*, 2006.

6 A. D. Edelstein, M. A. Tsuchida, N. Amodaj, H. Pinkard, R. D. Vale and N. Stuurman, *J. Biol. Methods*, 2014, **1**, 10.

7 J. A. Varela, M. Rodrigues, S. De, P. Flagmeier, S. Gandhi, C. M. Dobson, D. Klenerman and S. F. Lee, *Angew. Chemie Int. Ed.*, 2018, **57**, 4886–4890.
